# Embryonic development of fully biocompatible organic light-emitting diodes

**DOI:** 10.1101/2021.06.14.448347

**Authors:** Bruno F.E. Matarèse

**Affiliations:** Department of Physics, West Cambridge Site, Cambridge, CB3 0AS, UK; Department of Haematology, Cambridge biomedical Campus, Cambridge, CB2 0AW, UK

**Keywords:** *organic light-emitting diode OLED*, *light emitting polymer LEPs*, biocompatible, gold bioelectrode, bioelectronics, medical light source implant

## Abstract

Prototype fully biocompatible organic light-emitting diodes are investigated, with a view to creating a suitable and high-performance light source as a medical implant device. A selection of organic LED materials that have potential suitability for the biological environment are examined. First, the biocompatibility of selected OLED materials was evaluated by the study of cell adhesion and cytotoxicity of HeLa cells cultured on the candidate materials. Thus it was possible to design a device structure composed entirely of biocompatible materials. Second, the characterization of the electroluminescence properties of the prototype OLED is shown and its limitation evaluated. Third, the aqueous stability of the fully biocompatible light source is examined. There is strong evidence that fully biocompatible and stable light-emitting implant devices can be easily constructed. This is the first time a fully biocompatible organic light-emitting diode, albeit embryonic, is reported, with the hope that it may lead to further research to optimize the device performance. Some suggestions on suitable device properties towards *in vivo* transition are provided.

## I. Introduction

Optical imaging or detection (e.g. biosensors), and opsin based neuromodulation (e.g. optogenetics) are the major techniques that have changed the practice of optics applied in biomedicine. [1] Great advances have been made in the development of implantable medical devices related to size shrinkage, material suitability, system delivery such as wireless communication, electrical power consumption, and battery power. Society that is living ever longer demands solutions for a large number of clinical applications with the goal of developing miniature devices that can be fully implanted such as the pacemaker, cochlear implant or smart prosthetics. Whilst important results have already been obtained in this relatively new field, a significant additional problem facing researchers is how to provide light to specific deep brain areas that are of interest to address major clinical issues. To date, this has been achieved using inorganic light-emitting diodes (LEDs) or lasers as hardware (e.g. integrated fiber-optic and solid-state light sources) as conventional light sources that are able to deliver light with a precise wavelength to specific areas of the body (e.g. deep brain area).[2] However, these light sources are not biocompatible and are excessively bulky, which limits their effectiveness as body implants.

In contrast, an organic light-emitting diode (OLED) is a light source that combines optical and electrical properties with the known advantages of customized materials to provide appropriate color tunability, lightness, and low-cost solution processing.[3] The intrinsic multi-functionality of organic materials, such as light-emitting polymers, can be exploited to generate electroluminescence. Some of these light-emitting materials have been shown to be biocompatible with a wide range of biomedical uses [3]–[5]. In addition, mechanical flexibility combined with the high elastic modulus and biological inertness of carbon-based polymers and nanomaterials confer major advantages for use in non-conformal body cavities

OLEDs use organic semiconductors based on thin-film conjugated molecules or polymers as the active layer materials and they act as cold (non-incandescent), stable, energy-efficient solid-state light sources. They have simple sandwich structures, formed in the simplest case by sequentially depositing onto a plastic substrate a first electrode, one or more layers of organic semiconductor, and a second electrode. The electrodes and active layers are usually no more than a few hundred nanometres in height, so the total device thickness is determined in practice by the supporting substrate and any encapsulating layers used to protect against water and oxygen exposure, which together can be as thin as a few hundred microns. Engineering optoelectronic devices for operation in liquid environments are, however, a challenge that requires a more comprehensive understanding of the consequences that may arise in designing OLEDs for incorporation into living tissue.

This research is innovative in designing and engineering stable and biocompatible OLEDs for incorporation into living tissues. The aim is for organic LEDs not to alter cell viability, morphology and electrophysiological integrity/function. The principal challenge is ensuring they can operate in a highly saline and biologically active aqueous environment. Finding materials that are inherently stable in the environment in which they function is key to device optimization. In this paper, the proof-of-concept *in-vitro* cell-culture viability characterization confirms the feasibility of OLEDs to be fully biocompatible light sources. It is found that the active emitting layer, Super Yellow light-emitting polymer (LEP) Livilux PDY-132, is biocompatible and suitable for use in devices. A stable and biocompatible gold bioelectrode is used in this prototype device as the anode and cathode and was selected for its known suitability in medical implants. In order to help the injection of electrons from the gold bioelectrode used as the cathode, a thin layer of aluminum was placed between the gold and the light-emitting polymer. The conducting polymer poly(3,4-ethylenedioxy-thiophene: poly(styrene sulfonate) (PEDOT: PSS), is shown to not alter cell adhesion and is used in this prototype device as a biocompatible hole injection and transport layer. Here, the device performances with or without aluminum layers will be characterized to further evaluate device biocompatibility. The electroluminescence properties of embryonic light-emitting devices will be described below. The performance of the device will also be shown in cell-culture configuration. Some suggestions on the strategies towards *in vivo* transition are discussed.

## II. Preparation and characterisation

### A. OLED preparation

Five organic light-emitting configurations were fabricated. The bio-anode (configuration Fig. 2d with or without lithium fluoride LiF layer), the bio-cathode (configuration Fig. 2c with or without 5nm of Al layer), the fully-bio OLEDs on glass (configuration Fig. 2e) or substrate-free (configuration Fig. 2f) and the conventional OLEDs (configuration Fig. 2b with or without LiF layer). OLEDs were fabricated using gold (Au; WF ~5.1 eV) as a biocompatible anode (Fig. 2d-f), and for comparison, Indium tin-oxide (ITO; WF ~ 4.8 eV) (Fig. 2b-c). A biocompatible SU8 resign was used as an effective adhesion layer for gold bioelectrodes to the glass substrate. Semi-transparent gold bioelectrode (Au), 30 μm-thick, were fabricated following the previously described method [6]. ITO and SU8/gold-coated glass substrates were treated by O2 plasma for 10 minutes. PEDOT: PSS (Clevios CH4083, LumTech Taiwan) was spin-cast on top of the anodes under ambient conditions, forming a 40 nm-thick film. The 30 nm-thick emitting layers (EML) of Super Yellow SY-PPV (PDY-132 from Merk) in chlorobenzene were spin-coated on top of the PEDOT: PSS from pre-filtered precursors under ambient conditions. The as-cast samples were then transferred to a vacuum deposition system. Following active layers deposition, a bioelectrode made of Gold of 100 nm thick was subsequently deposited by thermal evaporation under a high vacuum (< 3×10-7 mbar). For some comparative devices, a thin layer of aluminum Al (5 nm) (configuration Fig. 2c, e-f) or lithium Fluoride LiF (0.8 nm) was placed between the cathode and the light-emitting polymer (configuration Fig. 2b&d). The precursor solutions of fluorescent polymers (SY-PPV) were prepared by dissolving 6 mg material in 1mL Chlorobenzene and were spin-coated for 30 s at 1200 RPM on top of PEDOT:PSS. After deposition of the OLED stack, some devices were immediately protected by an upper biocompatible polystyrene PS barrier as bio-encapsulant (fig.2. f) to evaluate the device *in-vitro* immersion lifetime.

### B. OLED characterization

The forward-viewing current-voltage–luminance characteristics of these OLED devices were measured using a Keithley 2400 source meter, Keithley 2000 multimeter, and a calibrated Si photodiode (from RS components), which was placed at a distance of 2 cm from the devices. External quantum efficiencies (EQE) were calculated from on-axis irradiance assuming a Lambertian emission profile and accounting for photodiode quantum efficiency across the electroluminescence spectrum. The electroluminescence spectra were obtained by a fiber spectrometer (Flame-S-VIS-NIR-ES, Ocean Optics). All the measurements were carried out at room temperature under ambient conditions. The stability of, namely, the bio-anode, bio-cathode, fully-bio (on glass or substrate-free) or conventional OLEDs were investigated by being immersed in *in-vitro* conditions for a prolonged period of time relevant to possible clinical use. The non-encapsulated or encapsulated by SU8/polystyrene OLEDs were placed in different plastic petri dishes filled with Dulbecco’s Modified Eagle’s Medium (DMEM) (Medium), kept in an incubator at 37.5 °C, 95% humidity, and 5% CO_2_. Finally, a control set of OLEDs was placed in an empty petri dish (air). The relative luminance of each OLEDs was measured in air at a fixed voltage corresponding to an initial luminance of 1 cd/m^2^ for various time-point during their *in-vitro* immersion.

### C. Cell culture preparation

Human HeLa cells, obtained from ATCC, were cultured at 2×106 /ml in Dulbecco’s Modified Eagle’s Medium (DMEM)– supplemented with 10% fetal calf serum, 2 mM L-glutamine, 100 ug/ml penicillin, and 100 ug/ml streptomycin (Life Technologies) directly on the surface of the glass, gold (Au), SY-PPV, PEDOT:PSS, and aluminum (Al) samples. PEDOT:PSS (Clevios CH4083, LumTech Taiwan) were cross-linked with glycidyloxypropyl) trimethoxysilane (GOPS) to avoid delamination or dissolution during the biocompatibility study. All samples were incubated for 48h and maintained at 37 °C in a humidified atmosphere containing 5% CO_2_.

### D. Biocompatibility assay

Pyknotic, Esterase-inactivity, and healthy cells were detected using the LIVE/DEAD fluorescence viability/cytotoxicity dyes (DRAQ7 ex/em: 633/675 nm (for Pyknotic); Annexin V pacific blue ex/em: 410/455 nm (for Esterase-inactivity) and Calcein UltraBlue AM ex/em: 359/458 nm for healthy), from Thermoficher. The combinations of Pyknotic/Healthy or Pyknotic/Esterase-inactivity were chosen in order to not overlap with SY-PPV photoluminescence. Radical oxidative stress production was detected using fluorescence TECAN Spark microplate reader at Ex/Em = 650/675 nm using the fluorescent cellular ROS Assay Kit (Deep Red) ab186029 from Abcam. The attached cells were rinsed twice with imaging solution (in mM: 140 NaCl, 2.5 KCl, 1 MgCl2, 1.8 CaCl2, 20 HEPES) and were then incubated for 4 min with the dyes. Cells were washed again with imaging solutions prior to image acquisition in the same solution. All products were used as received, gently hand-mixed, and briefly centrifuged. All measurements were conducted in triplicate and expressed as mean ± standard deviation (S.D.).

## III. RESULTS AND DISCUSSION

The cell adhesion and function compatibility of representative materials used in the field of plastic electronics such as polymeric encapsulator, light emitting polymer, charge injection and transport interlayers and metal electrodes were investigated. Polyethylenimine (PEI) has been previously shown to be cytotoxic [7]–[9] and was used here as a negative control while glass substrate was employed as a biocompatible positive control. First a simple adhesion assay of HeLa cell on materials investigated and their controls was carried out. Gold (Au), aluminum (Al), cross-linked PEDOT:PSS and glass displayed good cell adhesion. However, similar to PEI, the electroluminescent polymer, namely SY-PPV, displayed poor cell adhesion levels on its surface measured at 24 hours after cell plating. The next step was to enhance the adhesion performance of the materials by surface modification. It was found in previous work that surface treatment with the protein laminin, a principal component of a brain’s extracellular matrix, improves cell-adhesion onto hydrophobic light-emitting polymers.[3] To this end, a bio-functionalization of SY-PPV surface with laminin was carried out. An overnight incubation at 10 M/μ dramatically and significantly increased adhesion of all the investigated OLED materials (fig.1b). While promising, the adhesion capability of (HeLa) is not a reliable indication of cell viability. In order to assess more accurately the health of the cells, the viability/cytotoxicity assay was performed for all samples. As expected, the fluorescence assays carried out 48 hours after cell plating revealed poor cell adhesion on PEI where the cell viability remained just above 25%. Apart from the low cell attachment, DRAQ7 dye signal was strongly detected in cells grown on PEI, indicative of necrosis. By contrast, all the selected OLED materials evaluated presented levels of cell viability and mortality comparable to that of the glass control (i.e. above 90%). No significant differences were found between glass, gold, SY-PPV, PEDOT:PSS and aluminum (fig.1a,c). Furthermore, to assess damage to HeLa cells attached to materials, reactive oxygen species (ROS) was used as a marker for membrane integrity. It measured the generation of extracellular ROS in HeLa cells. PEI is shown to manifest an increase in intracellular reactive oxygen species (ROS). (fig.1d) However, in contrast to the PEI control, membrane fragility or cytotoxicity caused by ROS was not observed for gold (Au), aluminum (Al), cross-linked PEDOT:PSS and light-emitting polymer Super Yellow SY-PPV. They maintained significantly low ROS levels similar to the value in cells that were coated onto the glass control (Fig. 1d). Taken together the data indicates that PEI is not a suitable electron injection material for implantable optoelectronic medical devices due to its cytotoxicity. However, the results suggest that gold (Au), aluminum (Al), cross-linked PEDOT:PSS and light-emitting Super Yellow polymer allow for good cell adhesion and maintenance of essential cell viability and integrity parameters.

**Fig. 1.**
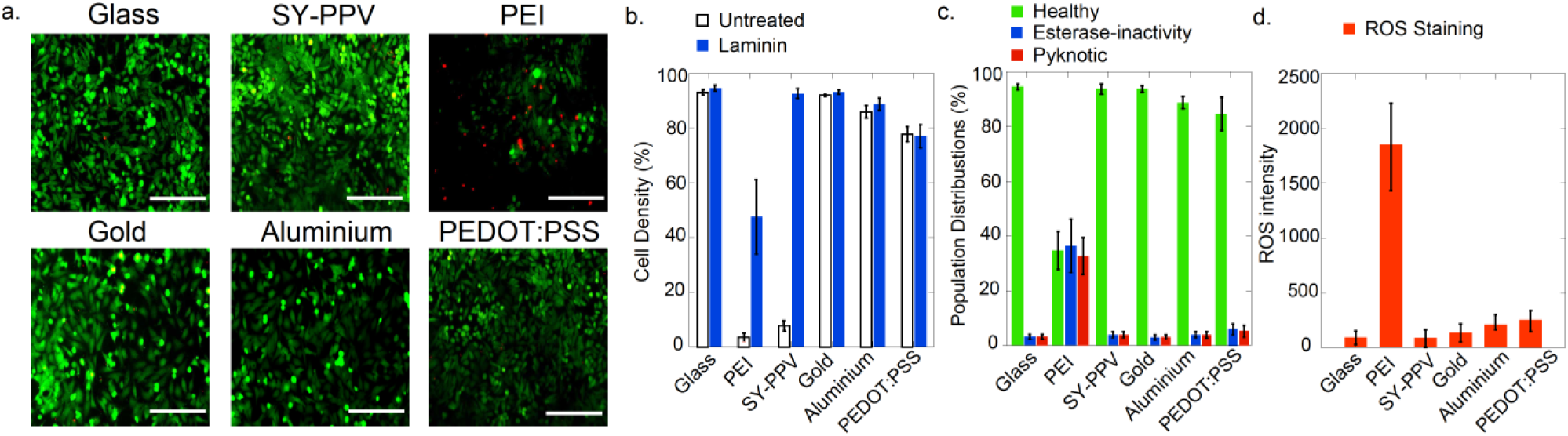
Viability of HeLa cell on glass, SY-PPV, Gold, Aluminum, PEDOT:PSS and PEI test-substrates. (a) Representative images acquired, with green regions signifying viable cells, red regions signifying pyknotic cells. Scale bare equal 100 um (b) Stacked bar charts derived from images, showing the population cell density for treated and untreated by Laminin. (c) Stacked bar charts derived from fluorescence images, showing the distribution of viable, pyknotic and esterase-free cells. (d) Stacked bar charts derived from fluorescence intensity, showing the ROS production. Error bars represent standard error of the mean of N=3 samples.

Following from the above it became possible to design a biocompatible prototype OLED device, which is suitable as an implantable optoelectronic medical device. On the cathode side of the device, the use of the gold biocathode demonstrated poor current density without any electroluminescence. This is due to poor alignment of the energy level between gold and the LUMO of SY-PPV (2.8 eV). However, the addition of a 5nm-thick layer of aluminum (configuration bio-cathode fig2.c & fully bio-OLEDs fig2.e-f with Al) generated electroluminescence via improved electron injection to SY-PPV. Bio-cathode & fully bio-OLEDs fabricated on glass substrate performed with high turn-on voltage of 21.4 V and 22.9 V; low current efficiency 0.06 Cd.A^−1^ and 0.03 Cd.A^−1^ and low external quantum efficiency EQE of 0.02% and 0.01%, respectively. Substrate-less fully bio-OLEDs electroluminescent characteristics were similar to the ones on glass. On the anode side, the use of the gold bioelectrode (configuration bio-anode & fully bio-OLEDs) showed lower current efficiency than devices using ITO anodes compared with the use the of same cathode configuration. For instance, conventional OLEDs (configuration fig2.b. using LiF as the electron injection layer) and bio-anode (configuration fig2.d. using LiF as the electron injection layer), have respectively turn-on voltage of 2.2 V and 3.1 V; low current Efficiency 5.17 Cd.A^−1^ and 2.44 Cd.A^−1^ and low external quantum efficiency EQE of 1.8% and 0.82%. The gold bioanode is actually less efficient partly because of its semi-transparency, with an average optical transmittance typically around 50% (compared to 89% for ITO) at its maximum peak approaching in wavelength the SY-PPV electroluminescence peak at 575 nm (fig2.g). Further optimization of the transmittance of bio-anode layer (e.g. using grid gold electrodes) or using other known biocompatible materials (e.g. graphene, carbon nanotubes CNT; highly conductive PEDOT:PSS, or hybrids), which are transparent to broader wavelengths, may improve fully biocompatible OLED performance. This would allow using blue-shifted or red-shifted biocompatible electroluminescent materials alternative to polymer Super Yellow. Alternatively, there is a need to identify bio-derived and/or biodegradable electroluminescent materials for biomaterial-based implantable photonic devices.

**Fig. 2.**
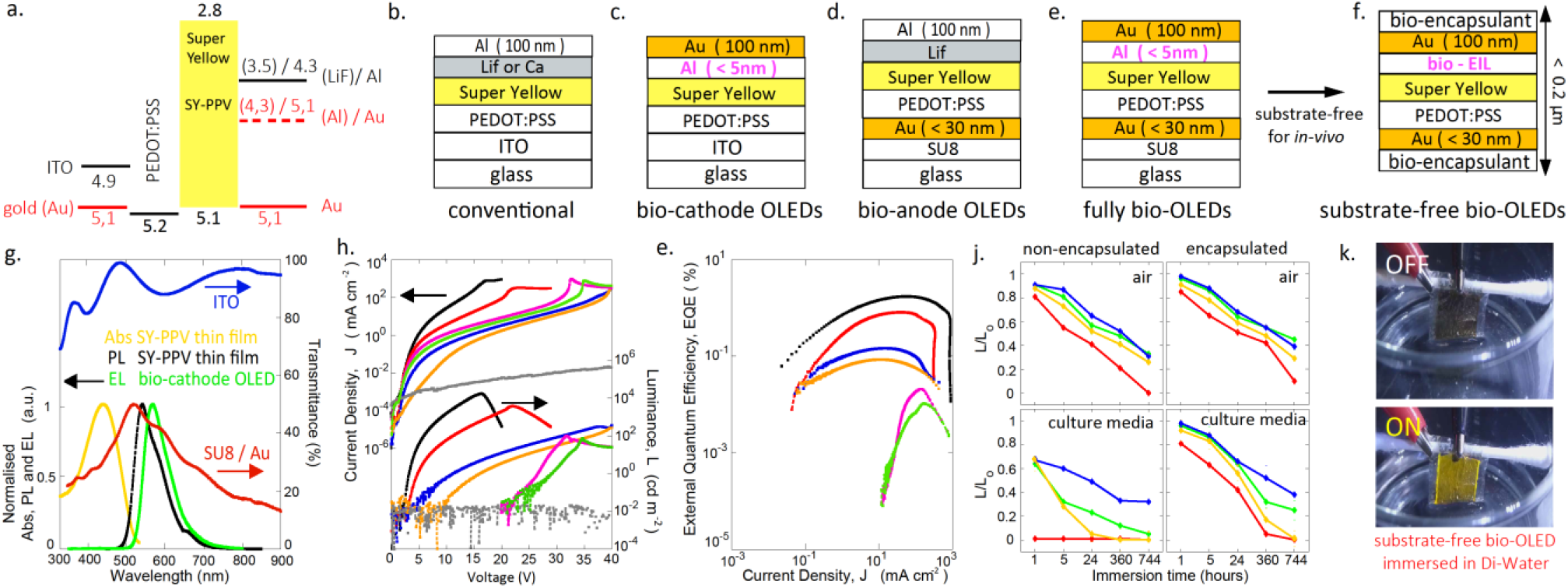
Electroluminescent characterization and stability of fully biocompatible OLED. (a) Energy level diagram. Schematic of stack architecture for (b) conventional; (c) bio-cathode; (d) bio-anode; (e) fully bio and (f) substrate-free fully bio OLEDs. (g) (right scale) Spectrum of absorbance (Abs) in yellow, photoluminescence (PL) in black for solid-state thin-film SY-PPV; electroluminescent spectrum in green for a bio-cathode OLED. (left scale) Transmittance spectrum ITO (in blue) and SU8/Au (in red). (h) Current density and Luminescence vs. voltage and (e) External quantum efficiency EQE v.s. Current density characteristics for ITO/SY-PPV/ LiF/Al (100nm) as Conventional (in black) ; ITO/ SY-PPV/AL(100nm) as Conventional without LiF (in bleu) ; Au /SY-PPV/LiF/Al (100nm) as bio-anode with LiF (in red); Au /SY-PPV/ Al(100nm) as bio-anode without LiF (in Orange); ITO/SY-PPV/Au(100nm) as bio-cathode without Al (in Gray); ITO/SY-PPV/Al(5nm)/Au(100nm) as bio-cathode with Al (5nm) (in pink) and Au/SY-PPV/Al (5nm)/Au(100nm) as fully bio with Al (5nm) (in green). (j) Relative luminance stability of non-encapsulated (left) and encapsulated devices (right) for bio-cathode (in blue); bio-anode(in red); fully bio(in green) and substrate-free fully bio OLEDs (in yellow) immersed in air (top) and cell-culture media (bottom). (k) Photograph of fully bio OLEDs immersed in Di-Water.

These parameters indicate the need for further optimization of the bio-anodes and bio-cathode configurations as well as combined for fully biocompatible OLEDs to outperform conventional non-biocompatible devices. Here, gold bioelectrode used as a cathode (for bio-cathode & fully bio-OLED with Al of 5nm) increases the air and cell-culture stability compared to conventional devices using LiF/Al (fig 2.j and table 1).

**Table 1.** Summary of device performance for bio-anode; bio cathode and bio-OLED compared to conventional OLEDs.

| OLED configuration | Turn-on Voltage ($V_{on}$) | LE<sub>max</sub> (cd/A) | EQE<sub>max</sub> (%) | Half -life time ($t_{1/2}$)<sup>a</sup> |
| --- | --- | --- | --- | --- |
| conventional | 2.2 | 5.17 | 1.8 | 24 h |
| bio-cathode | 21.4 | 0.06 | 0.02 | 744 h |
| bio-anode | 9.1 | 0.25 | 0.09 | 24 h |
| fully bio | 22.9 | 0.03 | 0.01 | 360 h |
| substrate-free fully-bio | 23.4 | 0.029 | 0.01 | 360 h |

a.) Half -life time relative luminance ( $t_{1/2}$ for $L/L_0=0.5$ ) for encapsulated OLEDs immersed in cell-culture media

Device immersion of the fully bio-OLED in cell culture for two weeks did not significantly affect its luminescence, in contrast to the effect on conventional OLED when only immersed for 24 hours. Hole transport material such as PEDOT: PSS has proven electrical stability in cell culture media. However, it is subject to current delamination and requires to be cross-linked. Moreover, the biocompatibility of non-crosslinked PEDOT:PSS as well as the influence of cross-linking on device performance is still to be confirmed.

Prolonged immersion of optoelectronic thin-films and devices in cell culture medium also highlights the importance of cross-linked polymers, while maintaining their electrical, optical and morphological properties, to achieve *in-vivo* long-term stability. [3] Previous work studied hybrid organic/inorganic or mixing thermoset plastics and thermoplastic encapsulation structures that may protect the device against oxygenation and create appropriate water vapor barrier. [3], [10] Polymeric insulator materials (e.g. silk, cellulose or Parylene-C) play complementary and necessary roles at the interface of the OLED/bio environment ensuring the protection of biological cells/tissues function/integrity and the protection the device against oxidative or reductive processes. [3], [10] Here, polystyrene PS and SU8 used as bio-encapsulation dramatically increase the light-source stability in cell-culture compared to non-encapsulated OLEDs (fig. 2.j).

The critical challenge to achieve fully biocompatible efficient and stable OLEDs that are suitable for *in-vivo* use remains to identify high performing and water-stable electron injection and transport materials. Synthesizing novel organic semiconductors will give rise to further advances in mechanically and chemically stable OLEDs using cross-linked and biocompatible materials for interfacing with biology.

Organic LED devices have recently shown their utility for biological manipulation, using organic light-emitting polymers in devices for the optogenetic activation of primary neurons. [4] Additionally, OLED devices have recently been employed for intrinsic and extrinsic *in vivo* optical imaging of functional cortical architecture and dynamics. [11], [3] The present article is the first report of substrate-less organic light-emitting diode made entirely of biocompatible materials working in cell culture medium. This embryonic research demonstrating the stability and biocompatibility of suitable OLED materials will pave the way for the next generation of medical implantable light sources.

## IV. ACKNOWLEDGEMENT

This work was supported by CRUK awarded fund with grant code RBAG/368.

